# Functional localization of visual motion area FST in humans

**DOI:** 10.1101/2024.09.19.614014

**Authors:** Puti Wen, Rania Ezzo, Lowell W. Thompson, Ari Rosenberg, Michael S. Landy, Bas Rokers

## Abstract

The fundus of the superior temporal sulcus (FST) in macaques is implicated in the processing of complex motion signals, yet a human homolog remains elusive. Here we considered potential localizers and evaluated their effectiveness in delineating putative FST (pFST), from hMT and MST, two nearby motion-sensitive areas in humans. Nine healthy participants underwent scanning sessions with 2D and 3D motion localizers, as well as population receptive field (pRF) mapping. We observed consistent anterior and inferior activation relative to hMT and MST in response to stimuli that contained coherent 3D, but not 2D, motion. Motion opponency and myelination measures further validated the functional and structural distinction between pFST and hMT/MST. At the same time, standard pRF mapping techniques that reveal the visual field organization of hMT/MST proved suboptimal for delineating pFST. Our findings provide a robust framework for localizing pFST in humans, and underscore its distinct functional role in motion processing.

## Introduction

The posterior bank of the macaque superior temporal sulcus (STS) contains several visual motion areas. These include the middle temporal area (MT), the medial superior temporal area (MST), and the fundus of the superior temporal sulcus (FST). In humans, visual motion areas are located in the ascending limb of the inferior temporal sulcus (ITS), where homologs of MT and MST are found. The preservation of FST across a range of non-human primate species including macaques ^1,2^, squirrel monkeys, owl monkeys, marmosets, and galagos ^3^ suggests that a homolog could exist in humans as part of the motion-processing system. There are, however, no established methods to localize or verify the existence of a human homolog of FST. Therefore, the present study evaluates the effectiveness of potential functional localizers for human FST and evaluates the case for homology using independent measures including motion opponency, retinotopic organization, and myelination.

Macaque FST receives major direct projections from MT ^1–4^, a region with an established role in the analysis of visual motion ^5–8^. Although both MT and FST process motion, they play distinct roles, and there is growing evidence that a primary distinction could be related to processing complex motion signals that extrapolate beyond retinal signals ^9^. A monkey neuroimaging study showed that FST responded more strongly to 3D motion compared to MT ^10^. This was supported by a recent electrophysiology study, which found that 37% of FST neurons but only 8% of MT neurons are selective for 3D motion ^11^. Furthermore, while MT neurons suppress responses to opponent motion signals ^12,13^, FST neurons often respond similarly to a single direction of motion and stimuli containing opposite directions of motion ^14^. FST is also strongly activated by structure-from-motion stimuli, in which stimulus elements can move in opposite directions ^15,16^. Finally, the involvement of FST in processing looming objects and its role in predicting their impact ^17^ as well as its contribution to action-related visual processing ^18^ further differentiate it from neighboring areas.

Previous work established human homologs of MT and MST using 2D-motion localizers ^19,20^. Both areas also show adaptation to 3D-motion stimuli ^21^ and direction of motion can be decoded from them as well as areas more anterior ^22^. However, these studies primarily relied on stimuli that contained both 3D and 2D (retinal) motion signals. A prior result, utilizing a stimulus that specified 3D motion in the absence of coherent 2D (retinal) motion signals, identified a region anterior and ventral to human MT/MST ^23^, echoing the location of FST in monkey studies. We reasoned that, based on results in monkeys, the use of a 2D motion localizer that does not contain 3D motion, and a 3D motion localizer that does not contain 2D motion, may dissociate human FST from neighboring MT and MST.

Recent advances in the understanding of the functional roles of macaque FST present an opportunity to establish a method to reliably and accurately localize FST in humans. To evaluate the case for homology, we present evidence for a putative FST region in the human using several distinct functional and structural MRI metrics. Importantly, the metrics considered are grounded in the unique characteristics of monkey FST, including its anatomy, response properties, receptive field size, retinotopic organization, and myelination.

## 2. Results

### 2.1. Criteria for localizing putative FST and hMT/MST

In our analyses, we first used the results from the 2D- and 3D-motion localizer stimuli to delineate hMT/MST from pFST. We then assessed the plausibility of pFST being functionally distinct from hMT/MST by assessing the differential activation of pFST to opponent motion, distinct population receptive field properties, and estimations of myelin density.

Traditionally, functional localizers are used to elicit activity from a particular cortical region whose function is distinct from neighboring regions. For example, it is common to select voxels that have greater responses to moving than static dots to isolate the MT complex, which contains several motion-selective areas ^24^. However, a single functional localizer may activate multiple cortical areas (e.g., several voxels in the primary visual cortex in addition to the MT complex may respond more to moving than static dots). Conversely, a single cortical area may respond to various functional localizers or visual features (e.g., V1 shows selectivities for both orientation and motion). Our study adheres to assumptions rooted in prior literature (mainly the macaque literature) to localize pFST. The existence of non-visual and non-motion-responsive cells within FST ^25^ complicates the interpretation of fMRI response amplitudes to motion stimuli and it is not clear if human FST is activated by 2D-motion localizers. Consequently, we posit the following assumptions to guide our delineation.

The 2D-motion localizer (Figure 1A) is expected to consistently activate hMT/MST, with the potential to also engage pFST. This assumption is informed by the established response of all three areas to 2D-motion stimuli, albeit with a predisposition for hMT/MST activation. We anticipate that the peak activation elicited by the 2D-motion localizer will lie within hMT/MST rather than pFST, reflecting the primary association of 2D-motion processing with the former regions. Conversely, the 3D-motion localizer (Figure 1B) is likely to activate pFST, given its specialized role in processing 3D motion ^11^. Activation of hMT/MST by this localizer is possible but not guaranteed given the heterogeneous findings about 3D-motion processing in macaque MT ^11,26,27^. Thus, we leave open the possibility that the peak activation for 3D motion may occur in either region of interest (ROI).

**Figure 1.**
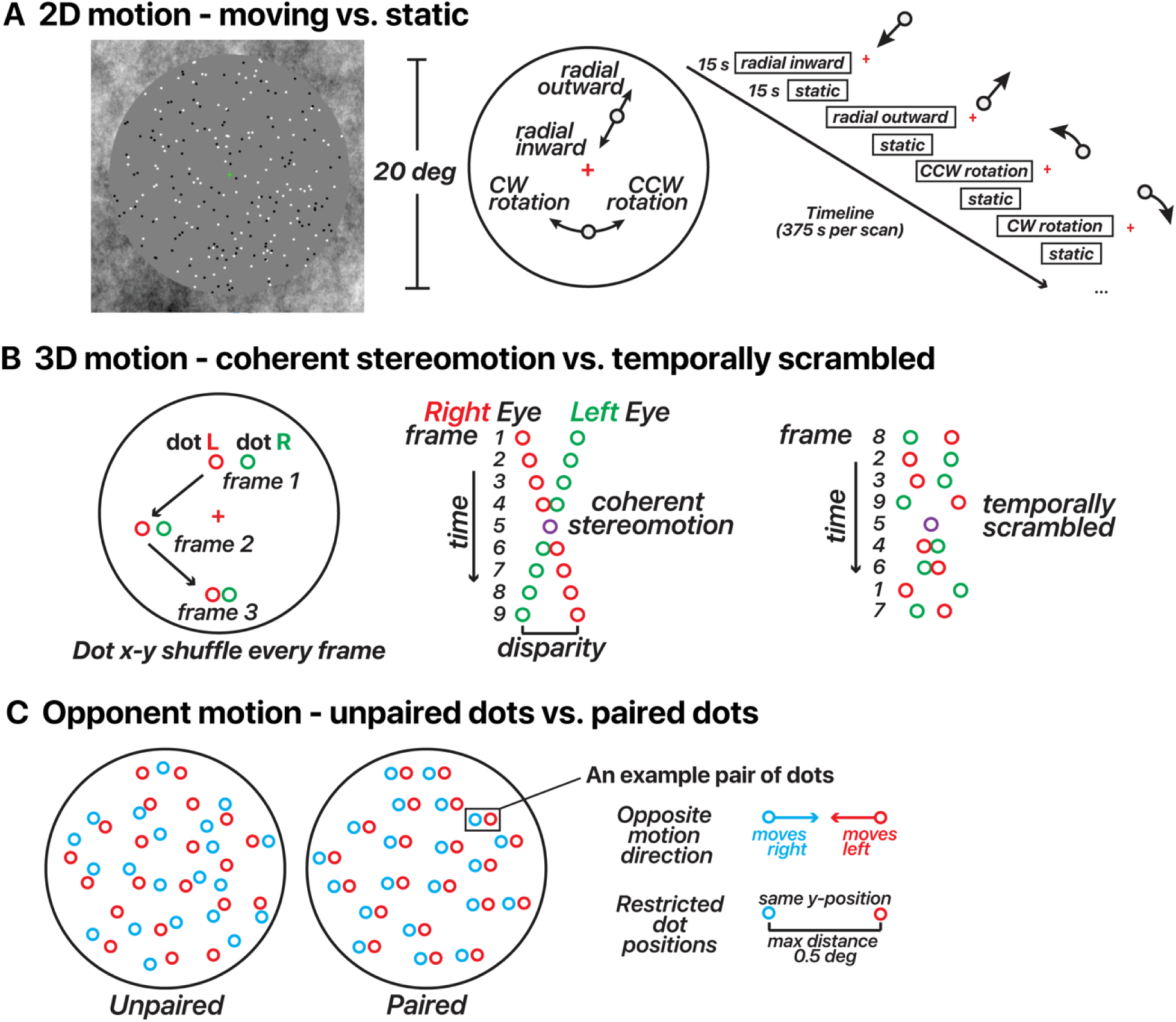
Motion stimulus design for 2D/3D-motion localizers and opponent motion. (**A**) 2D-motion localizer. Dots alternated from moving (inward, outward, clockwise, or counterclockwise) to static in 15 s blocks across the run. A fixation cross was located at the screen center. (**B**) 3D-motion localizer stimulus design (changing-disparity-defined stereomotion). Dots were binocularly presented. Red dots depict those presented to the right eye, and green dots depict those presented to the left eye. Red/green colors are used here for illustration. Dots displayed in blocks as either coherent stereomotion (perceived as 3D motion toward (1 s) – away (1 s) in a central disk and away (1 s) – toward (1 s) in a surrounding annulus) or temporally non-coherent (scrambled) motion in 10 s blocks across the run. In both conditions the dots’ *x*-*y* coordinates were shufled every stereo frame pair, the only difference was whether the disparity changed coherently or randomly (temporally scrambled order of stereo frame pairs). All dot pairs moved together in both conditions. (**C**) Opponent-motion stimulus design. Dots displayed in blocks as either unpaired (15 s) or paired (15 s). In both conditions, half of the dots moved rightward and half the dots moved leftward. For illustration, the dots were color coded in this panel by motion direction (red/blue); note that in the actual experiment all dots were white regardless of the motion direction and condition. In the unpaired condition, the dots’ positions were random. In the paired condition, the two dots within a given pair had the same *y*-coordinates and were never more than 0.5 deg apart.

These assumptions underpin our approach to delineating ROIs as illustrated in Figure 2. In some instances, we observed non-overlapping activations where 2D- and 3D-motion localizers elicited robust activation from distinct but neighboring areas (Figure 2C). In the majority of hemispheres, however, our results revealed a pattern where the 3D-motion localizer activated a larger area compared to the 2D-motion localizer. In these cases, the 2D localizer activated a subset of the 3D-activated region (Figure 2D). Following the above logic, the area activated by 2D and possibly 3D motion is identified as hMT/MST whereas the remaining nearby area activated by 3D but not 2D motion is identified as pFST.

**Figure 2.**
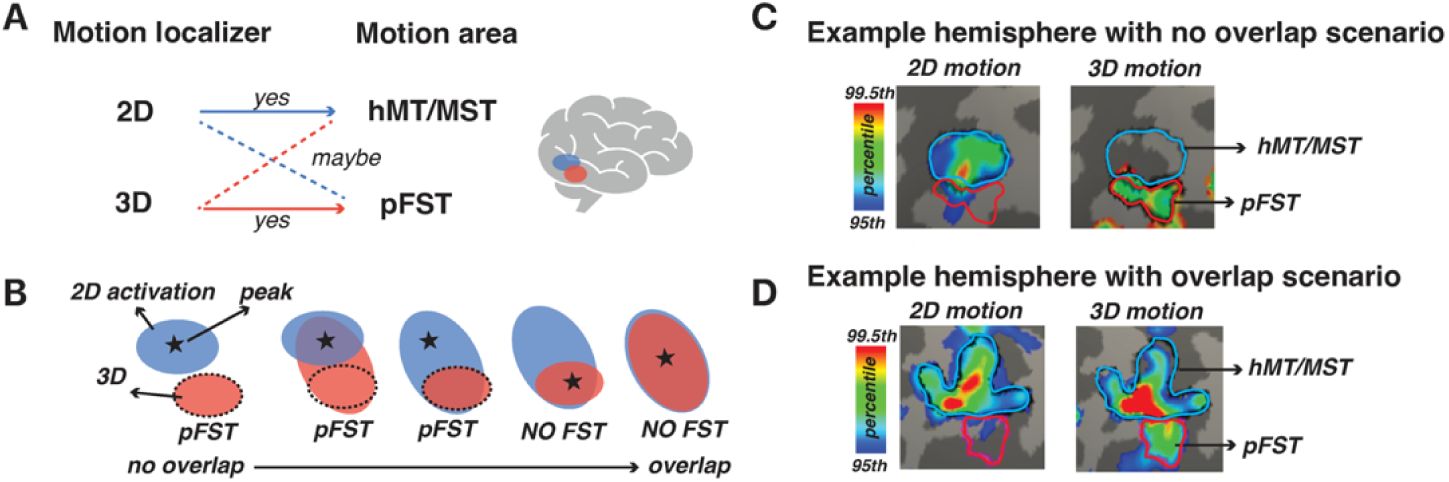
Delineating pFST. (**A**) Potential relationships between motion localizers and motion-processing areas. (**B**) drawing pFST in different scenarios based on the activation of 2D- and 3D-motion localizers. Blue patch represents selectivity to 2D motion, and the star symbol marks its peak activation. Red patches represent selectivity to 3D motion. The dashed circle marks the potential drawing of area pFST. (**C&D**) Example scenarios, zoomed in view on inflated surface, results thresholded at 95^th^ percentile. The blue outline represents hMT/MST and the red outline represents pFST.

### 2.2. Location of pFST across individual hemispheres

Figure 3 illustrates the location of pFST (colored red) in comparison to hMT/MST (colored blue) across individual hemispheres, presented on both white and pial surfaces as well as in a glass-brain view. We consistently localized pFST anterior and/or inferior to hMT/MST, a pattern that remains relatively consistent across individuals when viewed in the volume. Anatomically, and consistent with prior work ^28^, both pFST and hMT/MST were found along the ascending limb of the inferior temporal sulcus (ITS), near a sulcus that runs almost perpendicular to the ITS/STS, between the temporal and occipital lobes. While consistent in relative location to hMT/MST, the shape and size of pFST exhibited considerable variability between individuals, which is more noticeable when projected onto cortical surfaces. The largely symmetrical positioning of pFST across left and right hemispheres within individuals suggests some consistency in the neuroanatomical organization of motion processing.

**Figure 3.**
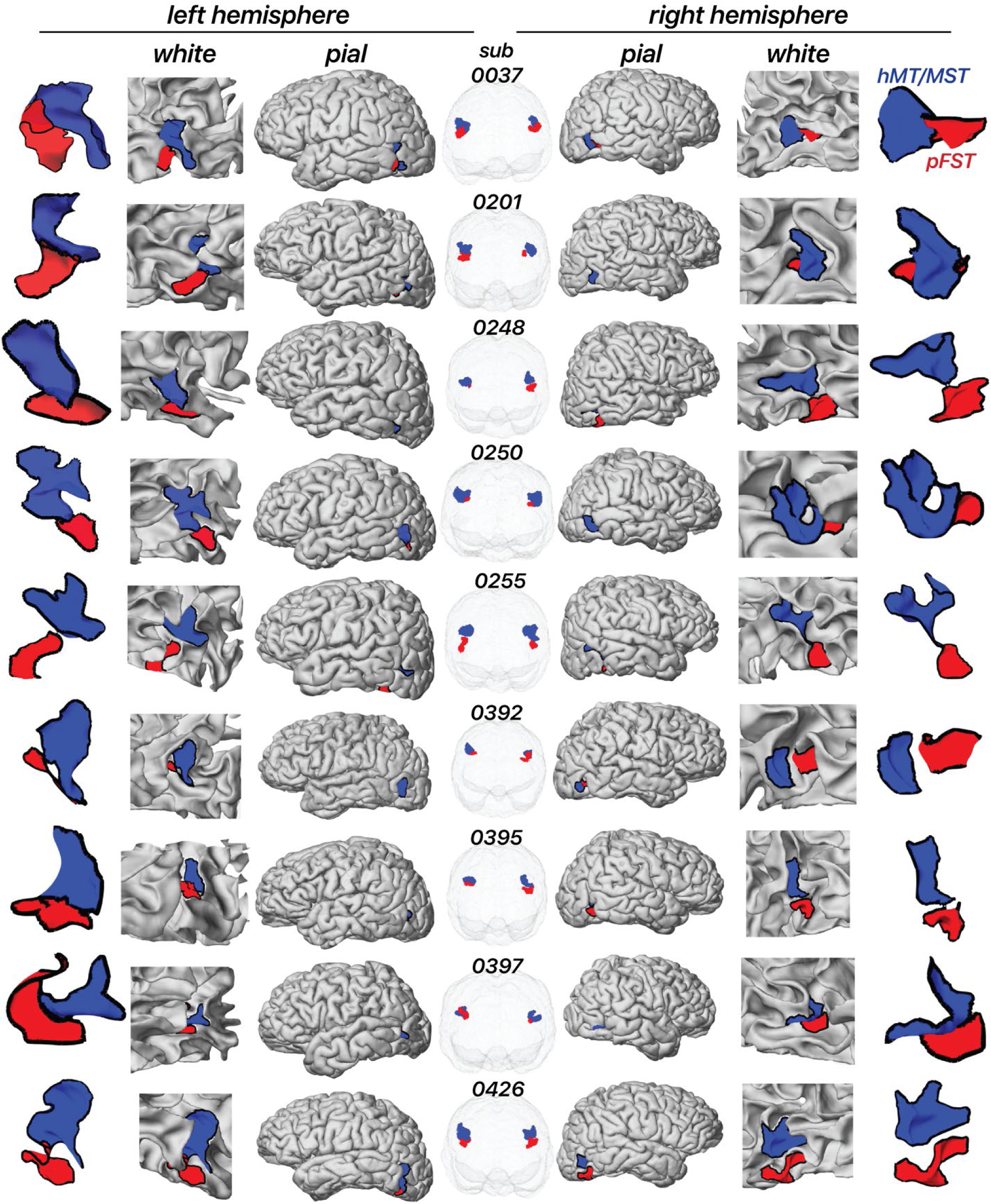
Location of area pFST and hMT/MST across individual hemispheres on the white and pial surface as well as in a glass-brain view. Each row represents a separate participant. The blue area represents hMT/MST and the red area represents pFST.

### 2.3. Evaluating complementary measures in a representative hemisphere

In each hemisphere, we defined the boundaries of hMT/MST and pFST based on differential responses to 2D- and 3D-motion localizers. Complementary measures were then evaluated to validate these boundaries. In monkeys, areas MT/MST are functionally characterized by significant signal suppression to opponent motion ^12,13^, and structurally characterized by high myelination ^4,29–31^. FST, on the other hand, shows little suppression to opponent motion ^14^ and is less myelinated in both monkeys ^3,32,33^ and humans ^34,35^.

In this section, we illustrate each of the analyses performed using an example hemisphere (see Section 2.5 for grouped results). In the example hemisphere in Figure 4, hMT/MST responded more strongly in the moving versus the static condition (*t*(295) = 32.49, *p* < 0.0001), as well as in the 3D stereomotion versus temporally scrambled control condition (*t*(295) = 16.42, *p* < 0.0001). Similarly, pFST also responded significantly more strongly in the 2D (*t*(94) = 13.42, *p* < 0.0001) compared to static, and in the 3D-motion conditions (*t*(94) = 25.53, *p* < 0.0001) compared to the temporally scrambled control. As expected, hMT/MST responded significantly stronger to 2D than to 3D motion (*t*(295) = 24.55, *p* < 0.0001). For pFST, the responses to 2D and 3D motion were not significantly different (*t*(94) = -1.7147, *p* = 0.0888).

**Figure 4.**
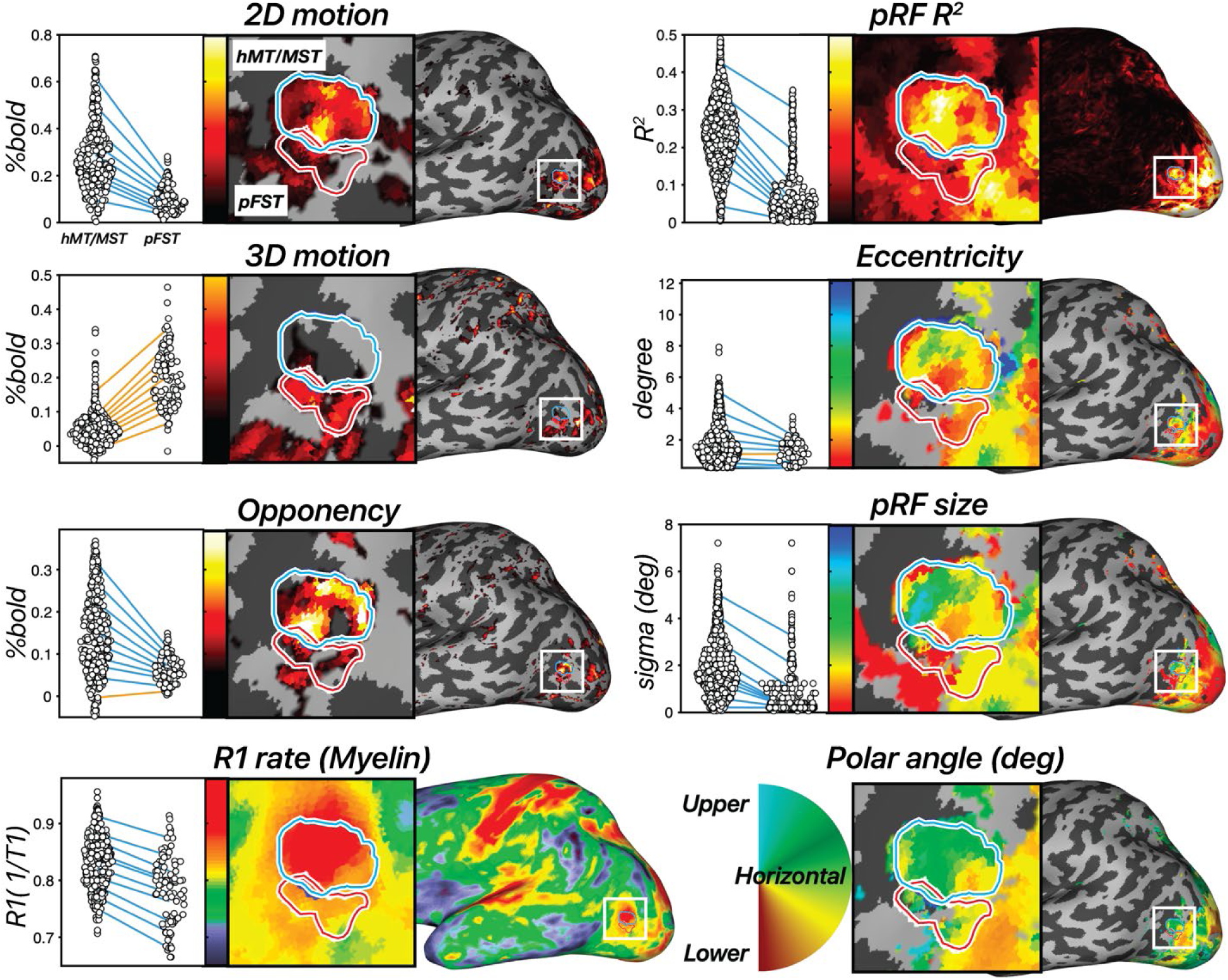
Results across 8 measures in an example hemisphere. hMT/MST is labeled with a blue outline and area pFST is labeled with a red outline. [left, first row] 2D-motion response (2D moving – static dots), [left, second row] 3D-motion response (coherent stereomotion – temporally scrambled dots w/disparity), and [left, third row] motion opponency (unpaired – paired moving dots) thresholded at the 90^th^ percentile. [left, fourth row] R1 rate (1/T1). Higher values are associated with greater myelin density. In the dot plot each dot represents a single vertex from the surface and each line connects each of the 10^th^ percentiles of the distribution across the two ROIs. Negative slopes (hMT/MST > pFST) are colored blue, and positive slopes are colored orange. [right, first to fourth rows] Estimated pRF parameters: variance explained (R^2^), eccentricity (deg), pRF size (deg), and polar angle (angle 0-360). All pRF results are thresholded at R^2^ > 10%.

We then used motion opponency as a complementary measure to verify the boundaries. The aim was to discern whether the region identified as pFST is indeed a separate area or simply an extension of the well-characterized hMT/MST area. In the example hemisphere, hMT/MST had significantly stronger signal suppression to opponent motion (Figure 1C) compared to pFST (*t*(389) = 8.29, *p* < 0.0001). This fits with the established understanding that areas like hMT, similar to their macaque counterparts, demonstrate a marked decrease in activity when presented with opponent-motion stimuli.

We also examined cortical myelination patterns using R1 rate, the inverse of the T1 relaxation time, as a proxy. We anticipated greater myelination in hMT/MST than pFST based on past literature ^3,32,34,35^. Average T1 relaxation time for gray matter in the human brain is about 1.331 s ^36^, which corresponds to a 0.751 s^−1^ R1 rate. Consistent with this, the median R1 rate across the whole hemisphere was 0.749 s^−1^ for this participant. The R1 rate for hMT/MST was 0.838 s^−1^ and for pFST was 0.792 s^−1^ with significantly greater values in hMT/MST than pFST (*t*(389) =7.69, *p* < 0.0001), suggesting greater myelination in hMT/MST. However, R1 rates in both hMT/MST and FST were significantly greater than the average cortical R1 rate (*t_hMT/MST_*(295) = 26.57, *p* < 0.0001; *t_pFST_*(94) = 4.76, *p* < 0.0001), suggesting that myelination in these motion-related areas may be greater than other cortical areas. This myelination contrast serves not as a localizing tool but as a confirmatory measure to validate the distinction between pFST and hMT/MST.

Additionally, 12 runs of the pRF-mapping stimulus were collected for each subject to compare the receptive-field properties between regions. Previously in the literature, two key points were relied upon in isolating FST. First, the presence of a significantly larger pRF size in FST compared to voxels in other areas at the same eccentricity. Second, a clear vertical meridian at the borders of FST with MST and V4t. There is a debate in the literature about the extent of retinotopic organization in FST. Some reported large receptive fields prioritizing central vision with no clear retinotopic organization in macaques ^4,25,30^, galagos ^3^, and Cebus apella monkeys ^32^. Others identified a complete visual-field representation of FST in macaques, including the lower vertical meridian separation from MST and the upper vertical meridian separation from V4t ^37^.

However, our pRF results did not ultimately prove useful in delineating pFST. Area pFST, in particular, showed low R^2^ values (median R^2^ = 11.17%), which compared unfavorably to hMT/MST (median R^2^ = 23.19%). This may be due to several factors. One possibility is that pFST has either extremely coarse or a complete lack of retinotopic organization. Another possibility is that pFST is retinotopically organized but cannot be topographically mapped using conventional methods due to constraints imposed by the limited field of view for visual stimulation within the scanner. Nevertheless, a lack of clear topology was a consistent observation across participants. We consequently ran several simulations to understand potential reasons for these outcomes.

### 2.4. Retinotopic mapping differentiates hMT/MST and pFST

Retinotopic mapping, and in particular population receptive-field mapping, can be used to localize and delineate areas along the visual processing hierarchy, including hMT and MST ^20,38,39^. However, this approach relies on the ability to reliably distinguish the receptive fields of different neural populations in response to localized stimuli across the visual field.

In monkey FST, receptive fields are substantially larger than in MT, with a radius that is larger by a factor of ∼1.5-2.2 ^11^. The visual field available for stimulation in a human MRI experiment is limited by the bore size of the scanner, typically limiting the field of view to the central 15-30 deg. In the experiments conducted here, stimuli were restricted to the central 24 deg. Practically speaking this meant the receptive-field size in pFST in humans would likely approach, or exceed the size of the available field of view, as defined by the stimulus aperture.

To empirically probe the efficacy of pRF mapping in pFST, we conducted retinotopic-mapping experiments in all of our observers. We subsequently fit two models to the data, a standard pRF model and a simple stimulus-contrast (ON/OFF) model. For the first model (pRF model), the responses depended on the visual field position of the bar stimulus relative to the pRF (peaking at timepoints when the stimulus maximally overlaps the pRF). For the second (stimulus-contrast) model, pRF responses were not specific to the stimulus position, but instead based on the total area of stimulus content, corresponding to the size of the aperture present on the display (peaking at timepoints when the stimulus/aperture covered the largest amount of the visual field).

We calculated the variance explained for each surface vertex using both models in V1, hMT/MST, and pFST (Figure 5). In V1, and to a lesser extent hMT/MST, variance explained was substantially greater for the pRF model than the stimulus-contrast model, suggesting that the spatial specificity of the neural population could be resolved within the stimulus aperture (diameter = 24.4 deg). In pFST on the other hand, variance explained was not substantially different between the two models. This suggested either that neurons in pFST are not retinotopically organized, or that the size of receptive fields within pFST approached or exceeded our stimulus aperture and therefore could not be estimated.

**Figure 5.**
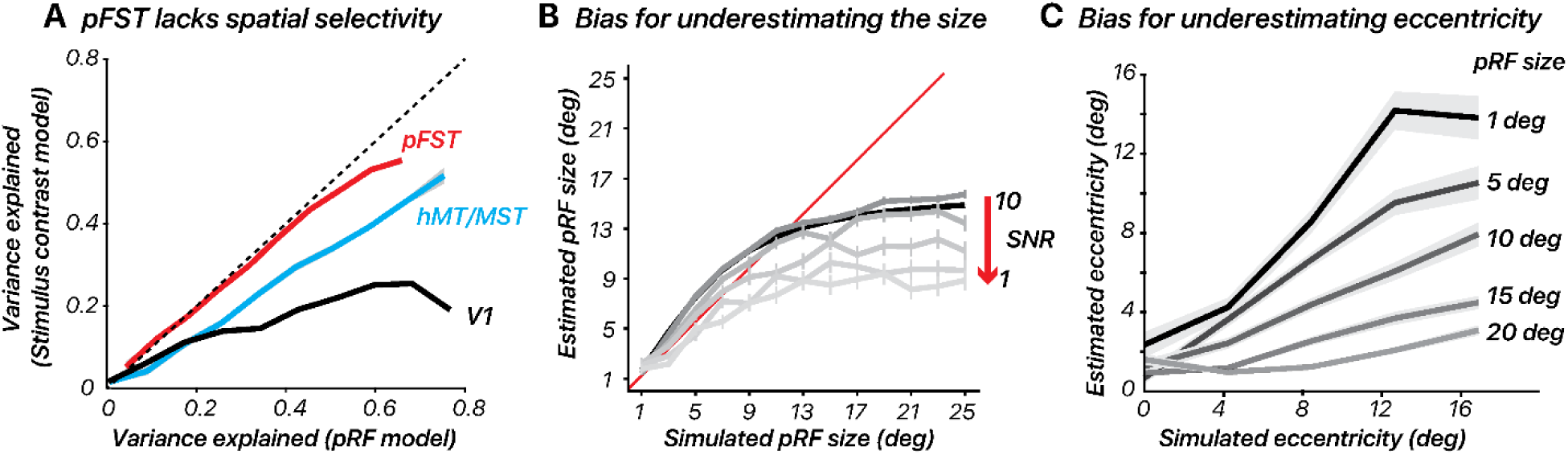
Retinotopic mapping differentiates hMT/MST and pFST. (**A**) Variance explained for one subject, using the pRF model versus stimulus contrast (ON/OFF) model in V1, hMT/MST, and pFST. (**B-C**) pRF size and eccentricity are underestimated for large pRFS, especially with low signal-to-noise (SNR). Error bars represent the standard error across bootstrap simulations.

Given that prior work proposed a pFST region with coarse topographic organization using fMRI ^39^, we investigated which factors could lead to a lack of clear topographic organization. To accomplish this, we assessed how noise levels influence estimated pRF size and eccentricity for areas with large receptive fields. We simulated BOLD time series for one of our retinotopic-mapping stimuli (the sweeping bar) and defined parameters for simulated pRF including eccentricity, polar angle, and size.

We simulated responses for these two models by iteratively increasing the receptive-field size and noise level. We expected these two factors to systematically result in poorer pRF estimates. Noise was defined as the standard deviation of Gaussian noise added to the modeled fMRI signals. By adjusting the noise standard deviation, the SNR was manipulated, simulating conditions ranging from low to high noise.

Consistent with our retinotopy estimates for pFST, the pRF simulations showed an underestimation of pRF size. This underestimation was more pronounced as noise levels increased (Figure 5B). We also evaluated how simulated pRF size influenced eccentricity estimates. We simulated pRFs with various sizes (1, 5, 10, 15, 20 degrees of visual angle) and center locations (0, 3, 6, 9, 12 degrees from the visual field center) for a group of 50 voxels. The fMRI response to each visual stimulus was calculated by convolving the stimulus profile with the hRF and adding noise. The pRF model was then applied to estimate the size and location of the pRF from the simulated fMRI data.

The results showed that estimating pRF eccentricity is inherently affected by the pRF size, with larger sizes significantly increasing the bias towards underestimating the eccentricity (Figure 5C). Taken together, these results confirm that retinotopic mapping is poorly suited to localizing and delineating visual areas whose receptive fields approach or exceed the size of the stimulus that can be presented. In our simulations, this led to results in which both pRF size and eccentricity were consistently underestimated. In cortical areas for which such concerns arise, simultaneously testing a simple stimulus-contrast model will be useful. If the variance explained does not improve meaningfully with a pRF model, this suggests that estimates of pRF size and eccentricity will be biased.

### 2.5. Triangulating human pFST: Group-level validation with motion opponency and myelination

While the current pRF methods may be suboptimal for pFST, the differences in pRF results between hMT/MST and pFST support the functional distinction between these regions. This distinction is further reinforced by two additional metrics: motion opponency and myelination (Figure 6). We identified greater motion opponency (activity for unpaired – paired motion) in hMT/MST compared to pFST across all hemispheres with a paired *t*-test (*t*(17) = 5.2307, *p* < 0.0001). In addition to the group-level results, individual one-tailed two-sample *t*-tests were carried out for each hemisphere, yielding significant results for stronger motion opponency in hMT/MST in 15 out of 18 hemispheres. The consistency of these results supports the functional distinction between hMT/MST and pFST. However, in three hemispheres (two from the same participant), there was no significant difference. This could suggest that, in a few instances, areas classified as pFST based on 3D-motion activation might partially overlap with hMT/MST.

**Figure 6.**
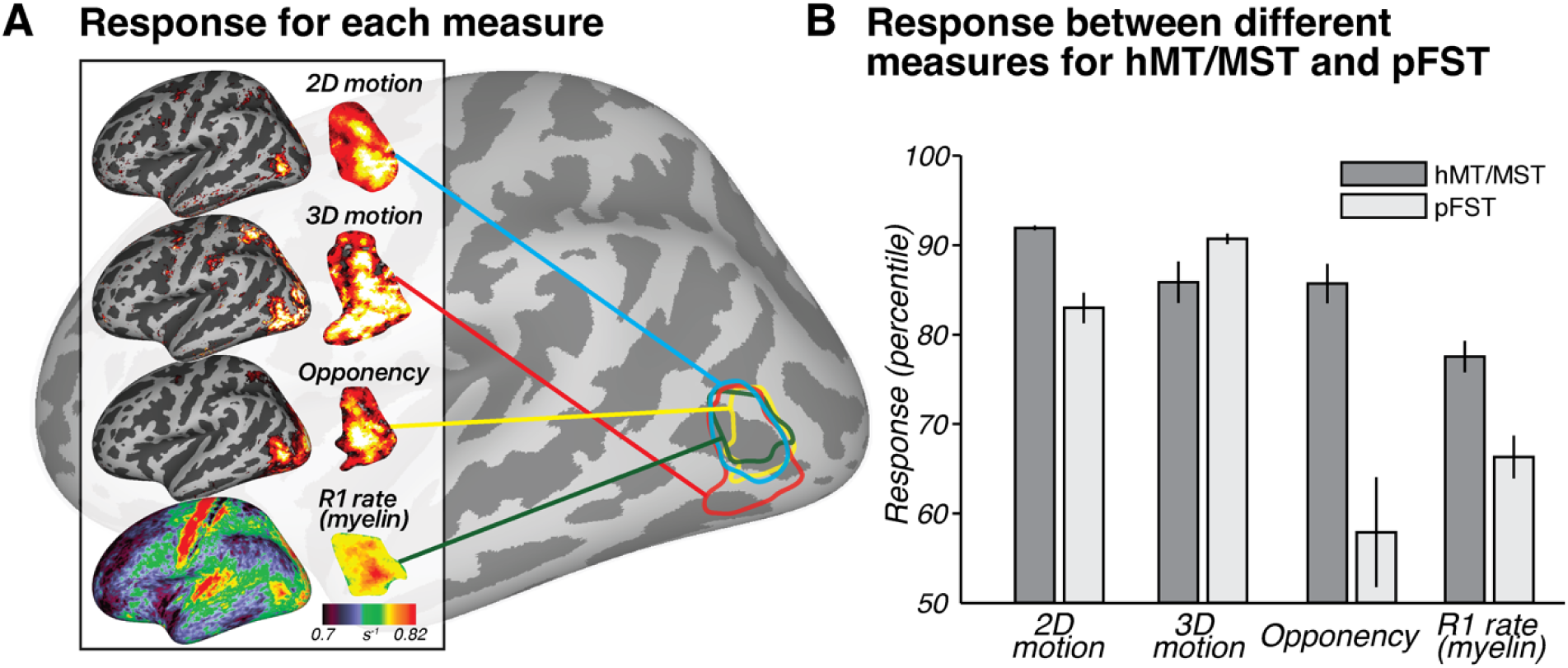
Group-level results. (**A**) Response for each measure across subjects (n = 9) to 2D motion (moving – static), 3D motion (coherent stereomotion – temporally scrambled), opponency (unpaired-– paired-dot motion), and R1 rate (myelin) visualized on an inflated left hemisphere on the fsaverage surface. For 2D, 3D, and opponency, color scale represents the percentage change in BOLD signal, thresholded from 90^th^ (red) to 99^th^ percentile (yellow). For R1 rate (myelin), color scale denotes variations in myelination ranging from 0.7 to 0.82 s^-1^. (**B**) Response between different measures for hMT/MST and pFST. Bars show mean and standard error across hemispheres (n = 18). Note that the values in **B** are plotted as percentiles for visualization purposes to emphasize the relative difference between hMT/MST and pFST. The statistical analyses reported in the text were conducted on the original values (BOLD signal change or R1 rate) without converting to percentiles.

A paired-sample *t*-test comparing myelination between hMT/MST and pFST across 18 hemispheres yielded a significant difference (*t*(17) = 5.1307, *p* < 0.0001). No significant differences in myelination were detected between hemispheres for hMT/MST (*t*(8) = -1.5324, *p* = 0.1640) or pFST (*t*(8) = -1.4400, *p* = 0.1878). The observed myelination patterns, in combination with the motion-opponency measures, support the conclusion that the areas we identified as pFST are functionally and structurally distinct from hMT/MST.

Interestingly, for one participant (sub-0392), both hemispheres displayed results in both the motion opponency and myelination measures that were atypical compared to the other participants. The hemispheric consistency of the results for that participant is consistent with individual cortical variability that may have reflected several identified factors. First, that participant’s average hMT/MST surface area across hemispheres was 181.53 mm², slightly more than half the average surface area of hMT/MST across the other eight participants (302.77 mm²). Second, the participant reported two preexisting medical conditions including regular ocular migraines and small lesions in V1. Because of these differences, we verified that excluding this participant had no effect on the statistical conclusions for any of the population results.

### 2.6. Individual variability compared to atlas-based location

We compared our functionally defined ROIs (Figure 7), based on responses to motion localizers, to those outlined by the Glasser et al. (2016) atlas. The Glasser atlas contains a comprehensive cortical parcellation, which is based on a combination of cortical architecture, function, connectivity, and topography. Unlike many other parcellation atlases, it contains an FST region. We quantified the agreement between the atlas-defined and functionally defined ROIs by quantifying their spatial overlap using each delineation method. We computed the percentage overlap per ROI by doubling the number of overlapping vertices and dividing by the total number of vertices in both the atlas-defined and functionally defined ROIs. The correspondence between functional and atlas-defined ROIs varied across subjects without a consistent deviation pattern. This variability was most pronounced for pFST, with hMT/MST generally showing a greater degree of overlap and smaller centroid distances, often less than 1 cm. Specifically, hMT/MST exhibited a median overlap of 43% and an average centroid distance of 0.68 mm in native space. For pFST, the median overlap was 11% and the centroid distance was 1.36 mm in native space. Functional pFST had little overlap with atlas hMT/MST (7%). Likewise, functional hMT/MST had little overlap with atlas FST (12%). Surface area of functional hMT/MST was 289.2 mm² on average across 18 hemispheres, about twice the size of functional pFST (153.4 mm²). For comparison, the atlas-defined surface areas were both larger than the functional ROIs, with hMT/MST at 379.4 mm² and FST at 301.7 mm².

**Figure 7.**
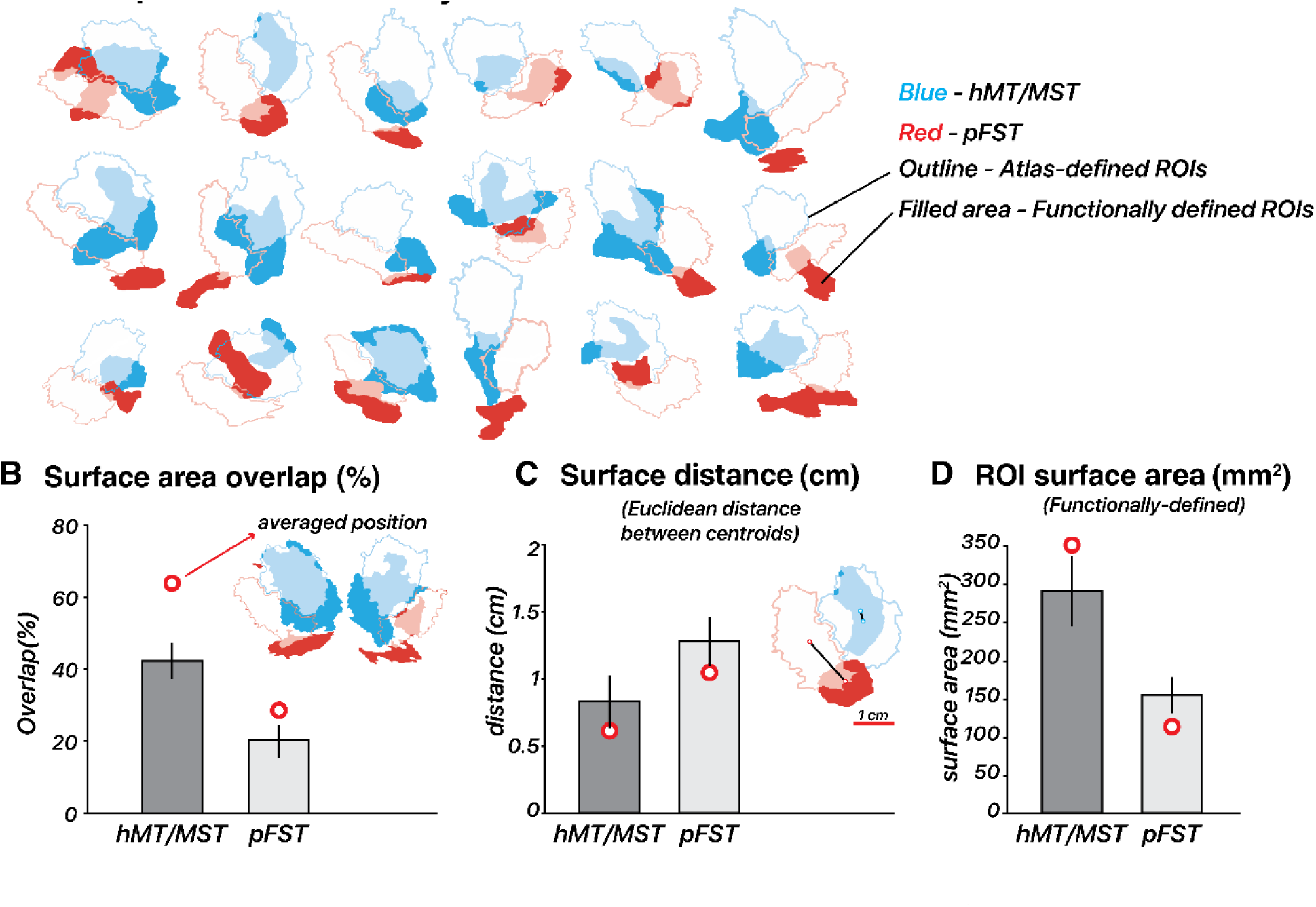
Functional vs. atlas-defined pFST. (**A**) Functional and atlas-defined ROIs in native inflated-surface space. Each cluster represents the overlap of regions delineated based on an atlas (Glasser et al., 2016) vs. functional localizer within a single hemisphere. Atlas-defined hMT and MST are combined into one ROI, which is comparable with functionally defined hMT/MST. Color-filled areas indicate vertices from ROIs manually drawn based on our localization criteria (2D- and 3D-motion functional localizers). Atlas-defined ROIs are shown as unfilled outlines. For both functional and atlas-defined ROIs, hMT/MST is filled in blue while pFST is filled in red. Filled areas with semi-transparent blue/red color indicate vertices that have consistent ROI labels for atlas- and functionally defined methods. (**B**) Surface area overlap (%): The bar plot shows the mean overlap percentage ± 1 standard error (black vertical line) across hemispheres in native space. Red circles mark the mean overlap value across subjects in fsaverage space. (**C**) Surface distance (cm): Distances in centimeters are calculated between the centroid coordinates of each ROI in inflated-surface space, with red circles marking the mean distance in fsaverage space. (**D**) Surface area (mm^2^): Mean surface area of functionally defined ROIs in inflated-surface space, with red circles marking the mean area of averaged functionally defined ROIs in fsaverage space.

To assess how well the atlas-defined ROIs aligned with the functionally defined ROIs’ average position across subjects, we transformed ROIs from all subjects into fsaverage space, and each vertex of the averaged surface was labeled based on majority agreement, reflecting the average functional position across subjects. In fsaverage space, the overlap percentage increased and the distances decreased for both ROIs. Functional and atlas-defined hMT/MST had 64% overlap, with a distance of 0.60 mm. Functional and atlas-defined pFST had 28% overlap, with a distance of 1.02 mm. In either native or averaged space, we observed significant variability across individual hemispheres. On average, atlas-based methods agreed well with the average functional data (red circle in Figure 7B), but when analyzing individual hemispheres, functionally defined hMT/MST and pFST deviated quite drastically from atlas-based parcellations (bars in Figure 7B). These deviations occurred for all subjects and could have a notable impact on statistical analyses given the relatively small number of voxels/vertices within these areas. Averaging across participants or relying solely on atlas-defined ROIs without accounting for such variability could obscure true anatomical and functional differences. This consideration is likely critical for motion-selective areas of the human inferior temporal cortex, including hMT/MST and FST.

## 3. Discussion

We demonstrated that FST’s unique functional properties set it apart from its neighboring motion-responsive areas located at the junction of the anterior occipital sulcus and the inferior temporal sulcus. Notably, our motion localizers revealed that pFST often responds less to 2D and more to 3D motion than other nearby cortical areas. The known sensitivity of FST to 3D motion in macaques both motivated and provided support that this area is indeed the human homolog. An additional functional criterion – reduced motion opponency ^14^ – validated this delineation method of pFST from other motion-selective areas. We summarized anatomical and structural characteristics that can further aid in identifying pFST, including its lower levels of myelin and anteroventral location relative to MT/MST.

Our delineation of FST was largely based on functional properties in the macaque, with some assumptions regarding the general proximity to MT/MST. Prior work with macaques supports the hypothesis that FST processes higher-order motion, including stereomotion ^9–11^ and structure-from-motion ^15,16^. FST is associated with the ventral pathway and thought to process the 3D motion of objects. This is consistent with FST’s strong connectivity to V4 and V4t, areas associated with the ventral pathway ^4^. In terms of the visual processing hierarchy, FST is considered to be downstream from MT/MST and to process more abstract forms of motion, and to be less selective for the direction of retinal motion than MT/MST ^30,40^. This macaque work is consistent with the functional responses and anatomical location of pFST that we highlight here.

Although not directly considered an FST homolog, a cortical area selective for stereomotion was previously identified in the human brain using a similar 3D-motion stimulus ^23^. The stereomotion region they located was adjacent to MT/MST, consistent with our findings, likely activating what we identify as pFST. However, their paradigm activated cortex that was on average anterior relative to MT/MST. We find similar anterior positions of pFST for some subjects (see Figure 3, e.g., subject 0037), but more often found pFST ventral (e.g., subject 0248) to other motion-selective areas. These differences may be attributed to individual variability in the precise cortical position of pFST. Differences in our stereomotion protocols may provide further explanation for the difference in average position. First, our coherent stereomotion condition contained two disparity-defined (toward/away) surfaces for which disparity increased (or decreased). We used disparity-defined surfaces based on evidence that FST responds strongly to surfaces such as motion-defined 3D shapes ^16^ and that selective processing of shapes/objects occurs ventral to MT/MST ^41^. Second, our control condition differed – our temporally scrambled condition contained several disparities (stimulus elements at different depths). Altogether, their study and ours both support the hypothesis that processing of stereomotion occurs adjacent to MT/MST. In addition, we both show that more complex visual-motion stimuli activate cortical areas that are: (1) not necessarily activated by 2D-motion localizers, and (2) associated with more “downstream” processing within the visual hierarchy.

### Population receptive field mapping

Retinotopic-mapping procedures are used to delineate several visual cortical regions based on their visual field maps, often by considering the eccentricity gradients and polar angle reversals that comprise a visual hemifield (or quarter field) representation ^42–44^. We found that the pFST boundaries were not reliably estimated using canonical retinotopic-mapping procedures. We conclude this for the following reasons. First, the parameter estimates of pRF eccentricity, polar angle, and size did not yield the common signatures of cortical retinotopic maps (e.g., full hemifield representations, or smooth gradients of size and/or visual-field position). Second, the explained variance tended to be quite low in these regions (less than 12%), making the pRF model estimates less reliable than for neighboring hMT/MST. Third, the pRFs were severely underestimated in size (< 0.5 deg) and biased toward the center of the visual field. Receptive fields in the macaque inferior temporal region are indeed known to overrepresent the fovea ^45^ but macaque FST in particular is known to have quite large RFs (∼8-35 deg) ^30^ at the eccentricities measured in this study. We therefore believe that the poor estimates result from noise and large RFs. Similar biases have been previously commented on ^46^. Additionally, cortical regions anterior and ventral to hMT have similar variance explained using a pRF model and an ON-OFF contrast model ^47^ suggesting a lack of (or greatly reduced) spatial selectivity in those areas.

But why is the signal from the mapping procedure so unreliable? Historically, macaque FST has been considered non-topographic ^48^ but a recent electrophysiology study showed a systematic change in size and eccentricity from posterior to anterior in FST ^11^ suggesting a coarse topographic organization. If this region is topographically organized in the human, the lack of such evidence in our data is likely due to the stimulus size. The very large receptive-field size of neurons in FST would exceed our 12.2 deg stimulus radius, thus requiring a larger stimulus display range to obtain topographic gradients of the visual field. Several groups have retinotopically mapped human MT and surrounding regions using fMRI ^20,38,39^. Kolster et al. used similar methods to identify a pFST region using retinotopic maps, largely relying on a shared foveal confluence between areas in the MT cluster. In this work, FST maps were very coarse and pRF size was smaller than in MT, potentially indicating a similar underestimation.

Surprisingly, despite using a smaller stimulus aperture (7.75 vs. 12.2 deg radius), they found gradual changes in eccentricity and polar-angle estimates. They highlight a slow duty-cycle (64 s or greater) of the retinotopic stimulus as optimal for these regions. Altogether, it is possible that pFST has retinotopic organization albeit with a much coarser spatial organization. To test this directly, the optimal stimuli would slowly sample space across a much larger window to obtain differential responses across neurons.

### Functional selectivity of FST & 3D-motion processing

Our localization of pFST should not be taken to indicate that this cortical area is exclusively dedicated to 3D-motion processing. There is speculation that the extrastriate body area (EBA) may overlap (at least partially) with FST. EBA is known to consist of three non-contiguous areas (LOS/MOG, MTG, and ITG) surrounding hMT+ that are selectively activated by images of limbs ^49^. The authors suggested that the limb-selective ITG likely overlaps with pFST. This is possible as human FST (localized by atlas) is activated by leg movements, unlike MT ^50^. This is in line with macaque FST, which includes non-visual motion-responsive cells ^25^. We can therefore expect that this area serves multiple functions across higher-level visual dimensions ^51^. As a separate but related point, the pFST region is not the only region activated by our 3D localizer. We have shown that MT/MST is somewhat responsive to 3D motion as well, suggesting that cortical areas that process 2D and 3D motion are not entirely distinct. The 2D localizer tends to activate less of the lateral occipital cortex than the 3D localizer, so we assume that cortical areas that are activated by 3D but not 2D motion signals must fall within FST.

### The human motion complex

Throughout this work, we refrained from making claims about whether FST lies within the human MT complex (hMT+). The MT region and its surrounding areas were initially termed a “complex” due to pending clarification, as it was understood that there were several motion-responsive areas forming a cluster in the macaque (including MT, MST, V4t, and FST) ^24^. However, in the context of most human fMRI studies, hMT+ is operationalized as the cortical region responsive to moving vs. static dots (2D-motion localizer), which surely includes MT and parts of MST, but may not include FST. To avoid giving too much credence to a term that is historically ambiguous, we instead focus on the ways that FST is functionally and structurally distinct from MT/MST. As delineation methods improve for motion-selective regions (e.g., based on functional specialization), the “complex” will either become a less useful term, or will need to be re-operationalized to make explicit what functions it includes. Our aim was also not to parcellate all possible areas of the MT complex. However, we wish to outline a potential subdivision that we did not address. In owl monkeys, FST is believed to include two distinct subareas, FSTd and FSTv ^33^. V4t, another region associated with the complex, was not considered in our study due to a lack of research specifically addressing its functional and anatomical properties in humans.

### From monkey to human

Our study is largely based on literature from monkey studies, the most prominent animal model used for insight into human visual processing. Although there are several commonalities among primate species regarding motion processing ^52^, there are also important differences between humans and monkeys ^15^. For example, motion area “MT” is in the middle temporal sulcus in the owl monkey ^33,53^, on the lateral surface of the marmoset ^54^, in the superior temporal sulcus of the macaque ^55,56^, and in the inferior temporal sulcus of humans ^28^. Additionally, these motion-selective regions exhibit significantly more positional variability in humans than other primates. This variability in humans partly results from greater variation in gyrification patterns, a co-varying factor with the location of the motion areas. Not only is there high variability in position, but also in the size and activation patterns in these motion areas. This variation occurs across individuals and across hemispheres within an individual.

The current study demonstrates that atlas-defined hMT/MST was generally consistent with the average functionally-defined hMT/MST – but this was not the case for pFST. When considering individual ROIs, these functionally defined areas were often misaligned with atlas-defined ROIs. The misalignment was often more extreme for pFST. We recommend that when conducting statistical analyses within pFST, and within most motion-selective regions surrounding MT, these analyses can be far more accurate when considering individual variability. It would be prudent to avoid the use of atlas-defined ROIs and caution should be used when averaging across hemispheres or individuals. Given the relatively small size of these ROIs (compared to V1), misalignment can result in a large proportion of cortex being omitted or mistakenly included for analysis.

### Technical limitations and methodological considerations

We do not have definitive parameters for the best 2D-motion localizer or opponent-motion stimuli. Our versions of these stimuli were designed as an initial proposal that can be built upon in future studies. The exact parameters used for each of the fMRI protocols were grounded in previous work; however, there is no certainty that these parameters (dot size, speeds, etc.) are optimal for activating FST. We limited the delineation of pFST based on the cortical responses to 2D- and 3D-motion localizers. Additional metrics including the protocols we used as validation criteria in this study (motion opponency, myelin, and anatomical priors such as size and position) could be used to refine the drawing of FST.

In conclusion, our study provides evidence for a distinct FST region in the human brain, characterized by unique functional and structural properties. FST likely plays a crucial role in the visual motion-processing network, particularly in integrating complex motion signals. It is involved in perceiving and interpreting 3D motion, which is vital for navigating and interacting with a dynamic environment. Additionally, FST’s role might extend beyond motion processing to include other sensory inputs, contributing to a more comprehensive understanding of spatial perception. Identifying the FST homolog in the human facilitates translational research between primate species and advances the mapping of sensory processing related to motion perception. Future research should aim to refine localization methods and explore the broader implications of FST in sensory-guided actions.

## Acknowledgements

NIH EY035005 (AR), NIH EY08266 (MS), ASPIRE, the technology program management pillar of Abu Dhabi’s Advanced Technology Research Council (ATRC), via the ASPIRE Precision Medicine Research Institute Abu Dhabi (ASPIREPMRIAD) award grant number VRI-20-10 (BR), and NYUAD Center for Brain and Health, funded by Tamkeen under NYU Abu Dhabi Research Institute grant CG012 (BR).

## Author contributions

Conceptualization: P.W., R.E., A.R., M.S.L, and B.R.; Methodology: P.W., L.W.T., and R.E.; Formal Analysis: P.W.; Investigation: P.W. and R.E.; Writing – Original Draft: P.W.; Writing – Review & Editing: P.W., R.E., L.W.T., A.R., M.S.L. and B.R.; Funding Acquisition: B.R., M.S.L., and A.R.; Supervision: B.R. and M.S.L.

## Materials and Methods

### Observers

Nine observers (four males, age 18 to 50 years) with normal or corrected-to-normal vision participated and provided written informed consent. All observers scored five or higher (70 s of arc or better) on the Randot Circles Stereotest (Stereo Optical Company, Chicago, IL). All observers participated in one scanning session for the functional localizers and two additional sessions for the population receptive field mapping. Each scanning session lasted 1.5 h. The experiments were approved by the University Committee on Activities Involving Human Subjects at New York University Abu Dhabi.

### Apparatus, display, and MRI data acquisition

We generated stimuli on a Macintosh computer using MATLAB 9.2 (The MathWorks, Natick, MA, USA) and the Psychophysics Toolbox extensions ^57–59^. Stimuli were presented using a ProPixx DLP LED projector (VPixx Technologies Inc., Saint-Bruno-de-Montarville, QC, Canada; refresh rate: 120 Hz, screen resolution: 1920 × 1080 pixels) with a rear projection screen (viewing distance: 88 cm; projected screen width: 38.5 cm) positioned at the back of the scanner. The display luminance was 107 cd/m² with a linearized lookup table. Stereoscopic presentation was achieved using a DepthQ Polarization Modulator from VPixx Technologies, placed in front of the ProPixx projector. Circular Polarizers from Edmund Optics were used as lenses, held in place by an MRIFocus lens holder from Cambridge Research Systems, mounted on a 64-channel head coil.

MRI data were acquired on a Siemens Prisma 3T full-body MRI scanner (Siemens Medical Solutions, Erlangen, Germany) using a 64-channel head coil. For each observer, a T1-weighted anatomical scan was acquired (TR: 2400 ms; TE: 2.22; flip angle: 8°; 0.8 mm isotropic voxels). This anatomical volume was used for white/gray matter segmentation and co-registration with the functional scans. T2^∗^-weighted functional scans were acquired using an echo-planar imaging (EPI) sequence (TR: 1000 ms; TE: 37 ms; flip angle: 68°, multiband factor: 6; matrix size: 104 × 104, 2 mm isotropic voxels; 72 slices).

We also collected MP2RAGE sequences to obtain additional T1-weighted images (TR: 5000 ms; TE: 2.98 ms; TI1: 700 ms; TI2: 2500 ms; flip angle 1: 4°; flip angle 2: 5°; 176 slices per slab; 1 mm thickness; echo spacing: 7.14 ms; slice partial Fourier: off). These images were merged to create a uniform T1-weighted image (UNI), from which T1 maps were estimated using qMRLab ^60^. The longitudinal relaxation rate (R1) was calculated as R1 = 1/T1 from the T1 maps and used to approximate myelin density.

### 2D-motion localizer

The 2D-motion localizer stimulus consisted of 250 black and white dots presented within a 10 deg radius circular aperture (Figure 1A*, see also Supplementary Video A*). Each dot was 0.2 deg in diameter and was presented for a limited lifetime of 0.5 s before reappearing in a random location. The background within the aperture was gray, and the area outside the aperture contained 1/*f* noise to aid fixation and vergence. The stimuli were presented using a block design with dots either static or moving (motion directions: radial inward, radial outward, counterclockwise rotation, and clockwise rotation). For the blocks containing moving dots, the velocity of each dot depended on eccentricity. For radial motion, the dot velocity increased as a function of the square root of eccentricity (maximum possible speed of 12 deg/s). For the rotational motion, we used one-eighth power scaling, rather than customary square root scaling to ensure reasonable movement across the stimulus aperture — square root scaling resulted in imperceptibly slow motion near the fovea. All four motion conditions had the same maximum speed of 12 deg/s. Each motion direction was repeated 3 times lasting 6 minutes per run and ended with 15 s of blank screen. Participants performed a color change-detection task to ensure fixation throughout each scan. A minimum of two scans were collected per participant.

### 3D-motion localizer

To attempt to delineate FST from other nearby motion-sensitive areas, we designed a stimulus that contained coherent 3D, but not 2D, motion signals (Figure 1B*, see also Supplementary Video B*). A similar stimulus design was used previously by Likova and Tyler (2007). There were 100 black and white dots presented within a grey aperture and 1/*f* background. The stimulus consisted of dynamic random-dot stereograms. When a participant fused the images received by each eye, this resulted in a perceived Z-depth for each fused dot; the perceived depth per dot depended on the binocular disparity of the dot pair. To strengthen the 3D percept, we created two disparity-defined non-overlapping surfaces (a circle and a surrounding annulus). The dots within the central 6 deg eccentricity always moved in the opposite direction in depth compared to the dots beyond that eccentricity. Stimuli included either coherent stereomotion (3D-motion condition) or temporally scrambled disparities (control condition), which were displayed in a blocked design. In the coherent stereomotion condition, the two disparity-defined surfaces started at the near and far sides of the volume (± 18 arc min disparity), and were perceived as moving in opposite directions, toward and away from the observer, both reversing direction every 1 s. The perceived 3D motion in this condition depended on the temporal (framewise) coherence of the disparity changes across all stereo dot pairs. The temporally scrambled condition (control) had temporally shuffled frames, retaining the range of binocular disparities presented (i.e., containing the same static depth information). Each possible relative disparity between the two disparity-defined surfaces was presented every second in both the coherent-stereomotion and scrambled conditions. The two surfaces remained discernible for each stereo frame pair, but the coherent 3D stereomotion percept was eliminated across frames.

Critically, the coherent-stereomotion stimulus contained 3D-motion signals (perceived as toward/away) in the absence of (spatially coherent) 2D motion signals. Thus, the two conditions were indistinguishable based on the monocular images; when closing either eye, the participant would perceive dots moving randomly in the image plane. Stereo dot pairs shared the same luminance. The stimulus alternated between the coherent-stereomotion and temporally scrambled condition every 10 s. Participants reported changes of the shape of the central fixation marker (o vs +) to ensure fixation throughout the scan. Each scan contained 5 minutes of stimulus presentation (15 repetitions of each condition) followed by 15 s of blank screen at the end.

### Opponent motion

We adapted stimuli used previously by Qian and Andersen (1994) to identify cortical areas with reduced responses to opponent motion (Figure 1C*, see also Supplementary Video C*). In each scan we alternated a paired-dots and an unpaired-dots condition (Figure 1C). In both conditions, 150 dots moved leftward and 150 dots moved rightward (constant speed of 5 deg/s). Each condition always consisted of 300 white dots per frame. In the paired-dots condition, the distance between dots within a pair could reach a maximum distance of 0.5 deg (resulting in a lifetime of 0.1 s). The dots within a pair moved one time across each other in the *x*-coordinate direction during their given lifetime. The dots within a pair also had equal *y*-positions. For the unpaired dots condition, the *y*-positions were random. The stimulus alternated between paired and unpaired conditions in 15 s blocks across the scan, and the entire scan lasted 5 m and 15 s.

### Receptive field mapping

We conducted experiments to map population receptive fields (pRFs) in putative FST and compared them to other motion-selective regions with topographic organization. We collected data for retinotopy using a stimulus and procedure previously described in detail ^47^. The stimulus content included colorful objects displayed on a 1/*f* noise background. The stimulus was confined within an aperture mask that changed position throughout the run. The procedure used two different types of apertures and data using each type of aperture were collected as separate scans. The apertures in the first scan were slowly rotating (clockwise/counterclockwise) wedges and expanding/contracting rings. The second type of aperture was a translating bar. To better drive motion-sensitive areas, additional runs were collected with an adapted version displaying moving stimuli behind the same apertures. The motion was created by expanding and contracting the original images in 0.6 s cycles. Each scan lasted 5 minutes, and 12 scans were collected in total: 6 scans which were the same as used by Benson et al. (2018) and 6 scans of the adapted moving version. For these scans, the stimulus-aperture radius was increased to 12.2 deg.

### Pre-processing and statistical analysis

All scans were organized using the Brain Imaging Data Structure (BIDS) format ^61^. We then used the fMRIPrep pipeline (version 20.2.1) for motion correction, spatial normalization, and co-registration of functional and anatomical scans ^62^. This step also converted our data into the fsaverage and fsnative spaces, which represent the data on the surface of the cortex with units of vertices instead of voxels. All subsequent analyses were conducted on the surface data. We ran a general linear model for all localizer analyses. The regressors included the condition onsets convolved with a canonical hemodynamic response function (hRF), a constant, a linear drift, and six translational and rotational motion regressors derived from fMRIprep. The beta weights were estimated separately for each run and subsequently averaged. Specifically, to derive the response to 2D motion, we subtracted the beta weights for static stimuli from the beta weights for moving stimuli. Throughout the remainder of this paper, we refer to the response to 2D motion as this contrast. Similarly, to derive the response to 3D motion, we subtracted the beta weights for the temporally scrambled condition from the beta weights for the coherent-stereomotion condition. For motion opponency, we subtracted the beta weights for the paired-dots condition from the beta weights for the unpaired-dots condition. We defined the strength of motion opponency as the degree of decrease in response between the unpaired- and paired-dots conditions. The above analyses resulted in one beta weight per vertex per metric (2D, 3D, opponency).

### Data and code availability

- Data needed to reproduce the results reported in this paper will be made available in a public data repository prior to publication.
- Analysis code has been deposited on GitHub and will be made publicly available prior to publication.
- Any additional information required to reanalyze the data reported in this paper is available upon request.

